# The effect of environmental enrichment on behavioral variability depends on genotype, behavior, and type of enrichment

**DOI:** 10.1101/557181

**Authors:** Jamilla Akhund-Zade, Sandra Ho, Chelsea O’Leary, Benjamin de Bivort

**Affiliations:** Department of Organismic and Evolutionary Biology & Center for Brain Science, Harvard University, Cambridge, MA 02138

**Keywords:** enrichment, behavioral variability, intragenotypic variability, microenvironmental effects, *Drosophila melanogaster*, high-throughput assay

## Abstract

Non-genetic individuality in behavior, also termed intragenotypic variability, has been observed across many different organisms. A potential cause of intragenotypic variability is sensitivity to minute environmental differences during development, even as major environmental parameters are kept constant. Animal enrichment paradigms often include the addition of environmental diversity, whether in the form of social interaction, novel objects, or exploratory opportunities. Enrichment could plausibly affect intragenotypic variability in opposing ways: it could cause an increase in variability due to the increase in microenvironmental variation, or a decrease in variability due to elimination of aberrant behavior as animals are taken out of impoverished laboratory conditions. In order to test our hypothesis, we assayed five isogenic *Drosophila melanogaster* lines raised in control and mild enrichment conditions, and one isogenic line under both mild and intense enrichment conditions. We compared the mean and variability of six behavioral metrics between our enriched fly populations and the laboratory housing control. We found that enrichment often caused a small increase in variability across most of our behaviors, but that the ultimate effect of enrichment on both behavioral means and variabilities was highly dependent on genotype and its interaction with the particular enrichment treatment. Our results support previous work on enrichment that presents a highly variable picture of its effects on both behavior and physiology

## Introduction

Stable behavioral differences among conspecifics are seen in a wide array of species. These differences, caused (definitionally) by some combination of genetic and environmental factors, are commonly referred to as individuality (Dall et al., 2004; Sih et al., 2004; Wolf and Weissing, 2010). Yet, even after experimentally homogenizing genotype and environment, individuality still persists (Gärtner, 2012), often undiminished or even increased (Ayroles et al., 2015; Buchanan et al., 2015). Multiple studies across different organisms demonstrate non-genetic individuality, which we refer to as intragenotypic variability (fruit flies: (Ayroles et al., 2015; Kain et al., 2012); pea aphid: (Schuett et al., 2011); nematodes: (Stern et al., 2017); fish: (Bierbach et al., 2017); crayfish: (Vogt et al., 2008); mice: (Freund et al., 2013; Kurikawa et al., 2018)).

This intragenotypic variability may originate in sensitivity to stochastic microenvironmental effects that persist even when large-scale differences in environment across individuals are removed (Debat and David, 2001; Honegger and de Bivort, 2018; Willmore et al., 2007). Along these lines, environmental causes of phenotypic differences can be decomposed into deterministic (macro) and stochastic (internal or micro) aspects (Clarke, 1998; Willmore et al., 2007). Examples of macroenvironmental effects are different levels of fertilizer or different temperatures across treatments. Examples of microenvironmental effects include whether an individual animal ate more food in the morning or evening, or (in the case of flies) whether they pupated on the plastic vial or food media surface. Generally, microenvironmental effects exist within a treatment regime (Debat and David, 2001) and are hard to measure (Honegger and de Bivort, 2018). For individuals of the same genotype raised in a homogenous experimental environment, trait differences would be primarily due to microenvironmental effects, and the propensity to this variation is known as microenvironmental plasticity (Morgante et al., 2015).

Many studies have focused on characterizing the intragenotypic variability in morphological, physiological, and behavioral traits (Abley et al., 2016; Ayroles et al., 2015; Bierbach et al., 2017; Blasco et al., 2017; Dworkin, 2005; Freund et al., 2013; Jacqueline L Sztepanacz et al., 2017; Kain et al., 2012; Kain et al., 2015; Mellert et al., 2016; Morgante et al., 2015; Sørensen et al., 2015; Tonsor et al., 2013). In *Drosophila melanogaster*, intragenotypic variability was found in chill coma recovery time, starvation resistance, sternopleural bristles, wing traits, and neuronal morphology in the larval ventral nerve cord (Dworkin, 2005; Jacqueline L Sztepanacz et al., 2017; Mellert et al., 2016; Morgante et al., 2015; Sørensen et al., 2015). Morphological variations present in the ventral nerve cord and optic lobes are of particular note as they respectively correlate with the timing of flight initiation and visually-guided locomotor biases, providing a link between morphological and behavioral intragenotypic variability (Linneweber et al., 2019; Mellert et al., 2016). Our research has identified intragenotypic variability in isogenic lines of *D. melanogaster* for turning bias, phototaxis, and thermotaxis (Ayroles et al., 2015; Kain et al., 2012; Kain et al., 2015). Outside of flies, intragenotypic variability in behavior has been studied in inbred mice and clonal fish (Amazon molly), with mice showing variation in exploratory behavior and fish showing variation in activity (Bierbach et al., 2017; Freund et al., 2013). If these examples of intragenotypic variability have their basis in microenvironmental differences, it may be hard to attribute the behavioral outcomes of specific individuals to their micro-causal underpinnings. It is, however, possible to test whether changing in the degree of microenvironmental variation predicts changes in the amount of intragenotypic variability.

As most lab organisms are already raised in heavily standardized environments where microenvironmental variation is minimized, it is feasible to increase microenvironmental variation and examine the effects on behavior. “Enrichment” treatments include a variety of different modifications to regular laboratory housing, such as opportunities for exercise, novel object interaction, and socialization (van Praag et al., 2000). Enrichment may add microenvironmental variation to a particular treatment, potentially affecting both the mean and variance of phenotypic traits (Körholz et al., 2018). Typically, enrichment treatments are hypothesized to more closely match an animal’s natural habitat, increasing mean well-being and cognition (while perhaps increasing intragenotypic variability). For mice and rats, enrichment has been shown to enhance mean gliogenesis, neurogenesis, and synapse formation in the cortex, hippocampus, and cerebellum, leading to improved memory and cognition (Bruel-Jungerman et al., 2005; Garthe et al., 2016; Leger et al., 2015; Mohammed et al., 2002; van Praag et al., 2000). Mice from enriched environments showed less activation of the amygdala, faster habituation to the forced swim test, and increased exploration in the open field test (Konkle et al., 2010; Mohammed et al., 2002; Van de Weerd et al., 2002). Salmon living in enriched tanks were faster in escaping mazes and showed increased expression of a pro-neurogenic gene (Salvanes et al., 2013). There is conflicting evidence for the effect of enrichment on brain size in fish, with enrichment having no effect on three-spined sticklebacks, but decreasing brain size in eastern mosquitofish (Toli et al., 2017; Turschwell and White, 2016). Early studies in *D. melanogaster* have found that changing the social milieu affects the size of brain structures, with social isolation leading to decreased sizes and numbers of Kenyon cell fibers (Barth and Heisenberg, 1997; Heisenberg et al., 1995; Technau, 1984). Social isolation in *D. melanogaster* leads to faster cancer progression, suggesting that a stimulating social environment buffers against stresses (Dawson et al., 2018). In crickets, mushroom body neurogenesis is higher in enriched environments with complex visual, olfactory, and auditory stimuli as compared to impoverished environments (Scotto Lomassese et al., 2000). On the other hand, enriched olfactory environments did not change mushroom body calyx size or affect odor learning in *D. melanogaster* (Wang et al., 2018).

The effect of enrichment on trait variability has been primarily studied in mice and rats and focused on understanding whether enriched rearing and housing conditions would decrease the statistical power to detect treatment effects by increasing within-sample variance (Toth et al., 2011). The evidence presented from behavioral and physiological studies has been conflicting - studies have shown that enrichment can increase, decrease, or have no effect on variability depending on the trait in question (Freund et al., 2015; Toth et al., 2011; van de Weerd et al., 1994; Van de Weerd et al., 1997; Van de Weerd et al., 2002; Wolfer et al., 2004). A recent study by Körholz et al. chose to focus more directly on whether intragenotypic variability in behavior and brain plasticity is influenced by the diversity of experiences that results from an enriched environment. They found that enrichment increases variation in specific domains - mice from enriched environments showed higher variation in exploratory behavior (object interaction times, habituation, but not locomotion), adult neurogenesis, and motor cortex thickness (Körholz et al., 2018). They attributed this increase in variation directly to the diversity of experiences or diversity of microenvironments that individuals could explore in an enriched environment.

Given the conflicting evidence for the effects of enrichment on variability, we propose two hypotheses for how enrichment may influence variation in traits. The first hypothesis is that enrichment introduces microenvironmental differences which in turn increase trait differences through microenvironmental plasticity. Our second hypothesis is that by more closely matching natural conditions to which organisms are adapted, enrichment increases the robustness of development and somatic maintenance, with a corresponding reduction in variation (due to the removal of aberrant phenotypes that may appear in impoverished laboratory conditions). Even though they predict opposite outcomes, both of these hypotheses are intuitive, and have some support in the literature. We chose to test them by measuring intragenotypic variability in *D. melanogaster* under control and enriched treatments. This species is a good model system for this work because of the ease of rearing large experimental groups from isogenic lines, and its suitability for automated behavioral phenotyping. We measured a variety of behavioral metrics associated with spontaneous locomotion and phototaxis (Ayroles et al., 2015; Buchanan et al., 2015; Kain et al., 2012) in five isogenic lines and two enrichment treatments (Figure 1). We found that while enrichment often caused a small increase in intragenotypic variability, the predominant determinants of behavioral means and variabilities were genotype and its interaction with the particular enrichment treatment.

## Results

### Intragenotypic variability is evident in locomotor and phototactic behaviors

In order to measure intragenotypic variability, we employed two automated assays which rapidly collect many behavioral observations from many individual flies. With Y-maze arrays (Buchanan et al., 2015), we measured the left-vs-right free locomotion turning bias of individual flies. With FlyVac (Kain et al., 2012), we measured the locomotory response to light cues of agitated flies (“fast phototaxis” (Scott, 1943)). We have previously used both of these assays to detect genetic and neural circuit regulators of intragenotypic variability (Ayroles et al., 2015; Buchanan et al., 2015; Kain et al., 2012), and between them, we examined both spontaneous and stimulus-evoked behaviors. We first confirmed that intragenotypic variability was present in a standard lab wild-type strain, Canton-S, in left-right turn bias and light-choice probability (Fig. 1). Indeed, the observed distributions of these measures were significantly broader (*p*<10^−3^ by bootstrapping, *χ*^2^, and Kolmogorov-Smirnov tests) than expected under null models in which all individuals behaved identically, i.e., sequences of behavior drawn from identical distributions (see Methods).

**Figure 1.**
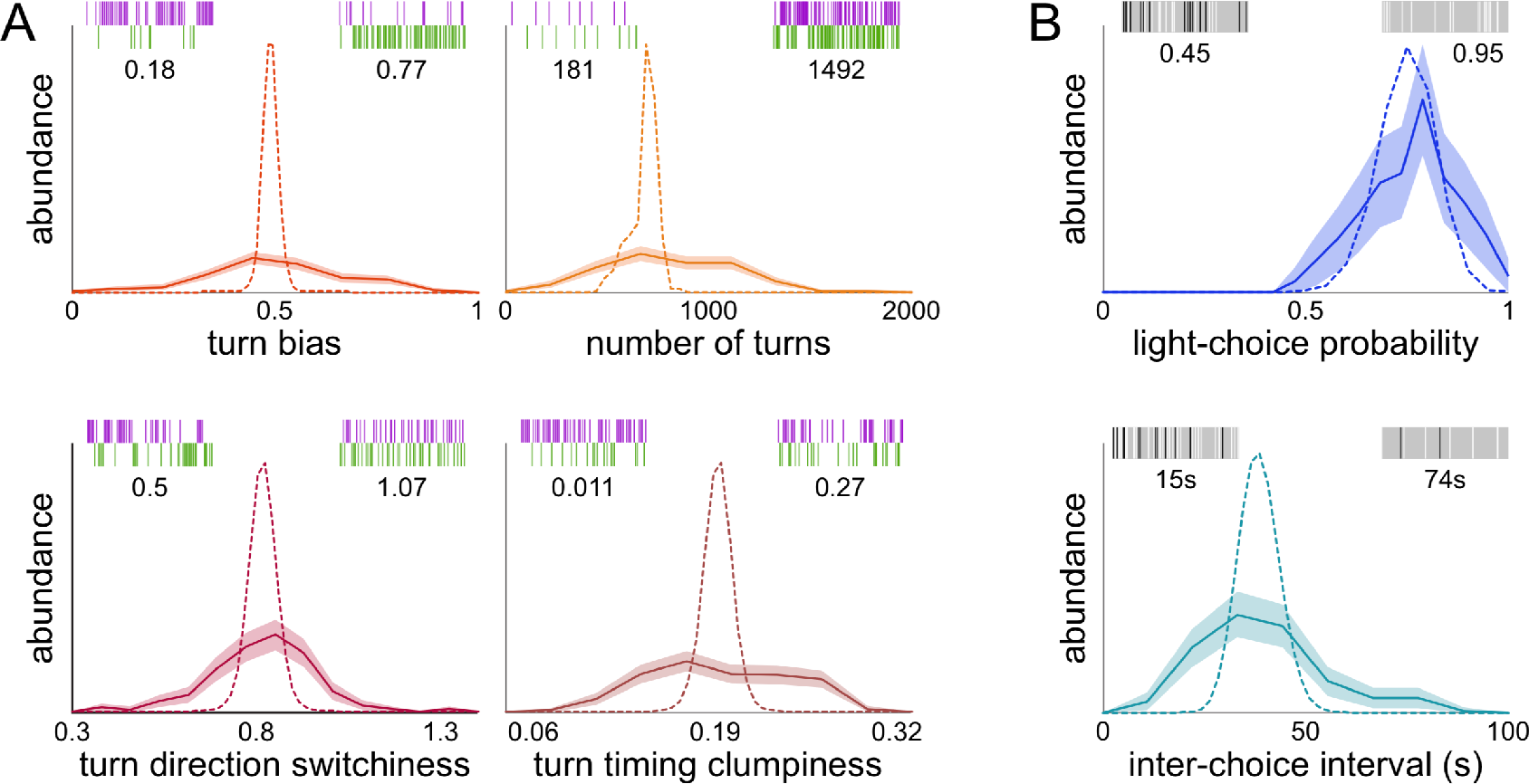
Observed and null hypothesis distributions of Y-maze and FlyVac behavioral measures for Canton-S (wild type) flies. Dotted lines represent the distributions expected under null hypotheses in which all individuals exhibit behaviors drawn from identical distributions. The solid line represents the observed distribution, with the shaded region representing +/− 1 standard error of the distribution, as estimated by bootstrap resampling. Insets show 10 minutes of representative data of the original behavior traces of extreme individuals and the corresponding value of that metric. A) Metrics from the Y-maze assay: turn bias is the fraction of turns made to the right, number of turns is the number of left-right choices made in the 2 hour test, turn direction switchiness is a turn bias-normalized measure of the mutual information between successive turns (for higher values, left turns are more predictive of subsequent left turns and vice versa), and turn timing clumpiness is a normalized measure of the irregularity of turns (the mean absolute deviation of the inter-turn intervals divided by the mean inter-turn interval). Purple ticks represent left turns and green represent right turns. B) Metrics from the FlyVac phototaxis assay: light-choice probability is the fraction of choices toward light, inter-choice interval is the mean time between choices. White ticks indicate a light choice, dark ticks show a dark choice, and shaded areas represent regions of time where no choice is made. 151 flies were analyzed for Y-maze behaviors, and 175 flies for FlyVac behaviors.

We next asked if there is evidence of intragenotypic variability in other measurements taken while measuring turn bias and phototactic preference in these assays. Beyond turn bias, we assessed 1) the number of turns completed by individual flies within the two hour trials, 2) flies’ tendencies to alternate between left and right turns successively (“switchiness”), and 3) the extent to which their turning events were clustered in time (“clumpiness”). As with turn bias, the observed distributions of these three measures were significantly broader than expected under null models in which all flies behaved identically (Fig. 1A). Using FlyVac data, we observed that the distribution across flies of the average interval between phototactic choices also was broader than expected if all flies were behaving identically (Fig. 1B). Thus, intragenotypic variability was evident in all six behavioral traits examined.

**Figure 2.**
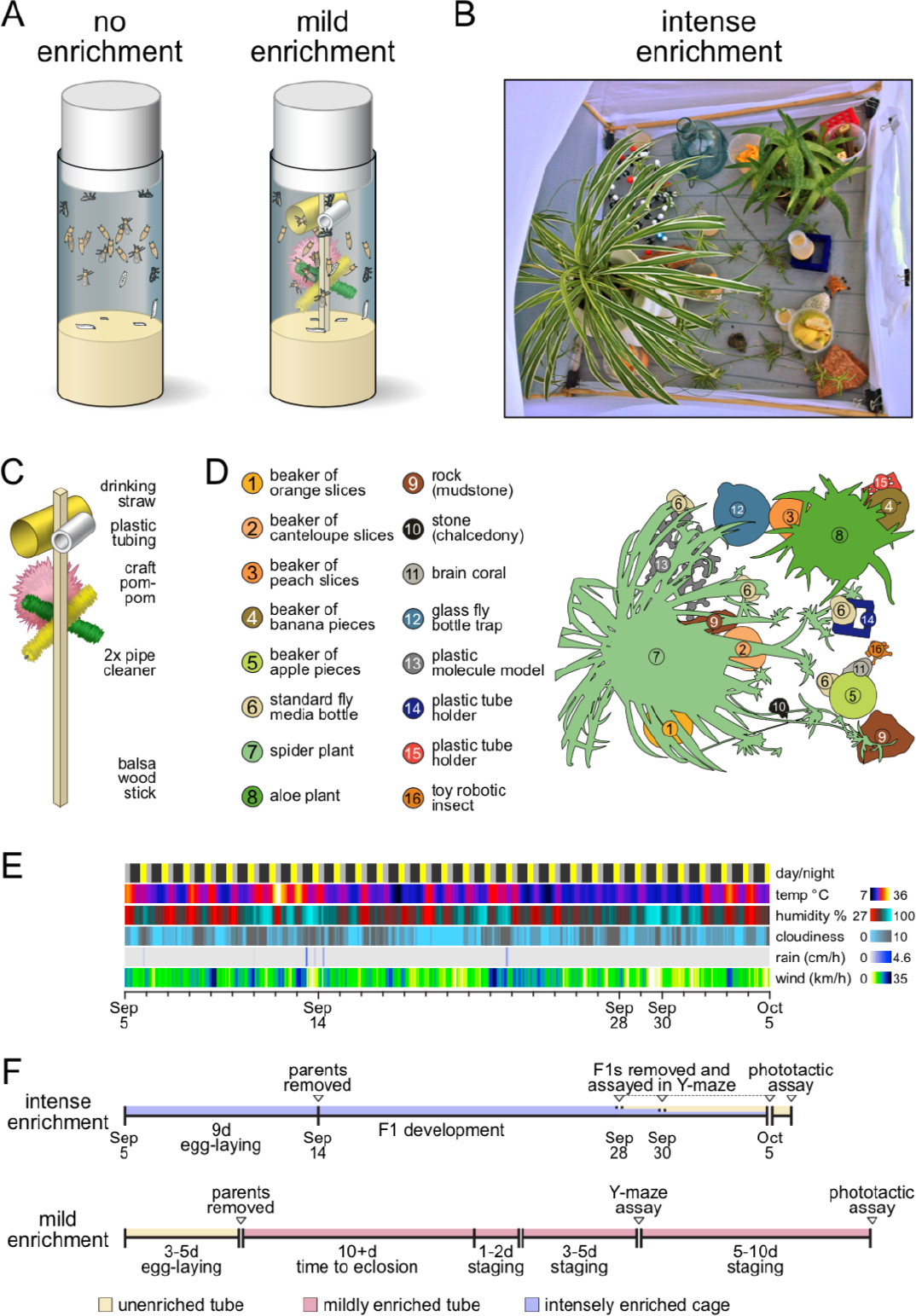
Illustration of enrichment paradigms used. A) Control vials vs. mild enrichment vials. B) Photo of the inside of the intense enrichment cage. C) Diagram of the mild enrichment ‘jungle gym’ components. D) Diagram of the intense enrichment cage components. E) Weather conditions that the intense enrichment cage was subject to for the experimental period. For the daylight timeline, yellow indicates potential direct sunlight on the cage, grey periods where the cage was shaded by our building, and black shows nighttime. The cloudiness timeline reflects the NOAA 10-point scale where zero is clear skies and ten is full cloud cover. F) Timelines of experiments showing the development, staging, and behavioral testing of the experimental animals for both mild and intense enrichment treatments. Each contiguous horizontal line indicates the time spent in a fresh container. The no enrichment control was the same as the mild enrichment, except all vials used were unenriched.

### Enrichment affects both mean and variability

We developed enrichment protocols that were either “mild” or “intense” (Fig. 2). For our mild treatment, we fabricated standardized “fly jungle gyms” that could be inserted into standard culture vials. These were made of small craft items: plastic tubing, pipe cleaners and fuzzy pompoms, mounted to a balsa wood stick (Fig. 2C). This enrichment treatment offered flies varied textural, luminance and color microenvironments. For our intense treatment, we constructed a 1m × 1m × 1m cage with insect netting as walls. Into this space we added 6 kinds of fly food (a variety of decomposing fruits as well as bottles of standard cornmeal media), houseplants, stones, varied plastic objects, etc. (Fig. 2D). The cage was placed outside on a deck where it experienced natural fluctuations in luminance, temperature, rainfall, wind, humidity, etc. during the course of our experiment (Fig. 2E). For both enrichment treatments and controls, we let a parental generation of flies lay eggs for 3-9 days, removed the parents, let the progeny develop and live for three or more days as adults before we removed them for behavioral testing (Fig. 2F).

To confirm that the enrichment treatments had an effect on our flies, we examined the mean values of our six behavioral phenotypes under each treatment. We used a Bayesian framework with a weakly informative prior to estimate the posterior distributions of the means and variances of each behavioral metric under the two enrichment treatments. We used the 99% highest density interval, also termed credible interval, (Kruschke, 2013; see Methods) to assess whether the posterior distributions of the means of each behavioral metric were different from each other. Intense enrichment caused strong decreases in the mean of number of turns and inter-choice interval, and a strong increase in the turn switchiness when compared to mild enrichment and the control (Fig. 3A). For these behaviors, intense enrichment had a larger effect on the mean than mild enrichment. Mild enrichment had a less pronounced effect on the mean number of turns and turn clumpiness. There was no apparent effect of enrichment on turn bias and light-choice probability, though the FlyVac assay has lower power than the Y-maze assay. The mean changes under enrichment indicated that our treatments had some effect on the flies, and could potentially alter intragenotypic variability as well.

We examined the effect of different types of enrichment on intragenotypic variability in our behavioral measures (Fig 3B). We chose to look at the coefficient of variation as our measure of intragenotypic variability in order to standardize it across multiple types of measures and control for mean effects (estimates of the posterior distributions of variance effects, not normalized by the treatment means, are included in Figs. S1,2). For nearly all behaviors, intense enrichment had a larger effect on variability than mild enrichment, but these effects where not all in the same direction. Intense enrichment decreased the variability of turn bias and turn direction switchiness, but increased the variability of number of turns and inter-choice interval, when compared to the control and mild enrichment treatments. Intense enrichment increased variability in clumpiness, though the effect was more pronounced upon comparison to mild enrichment as opposed to the control.

**Figure 3.**
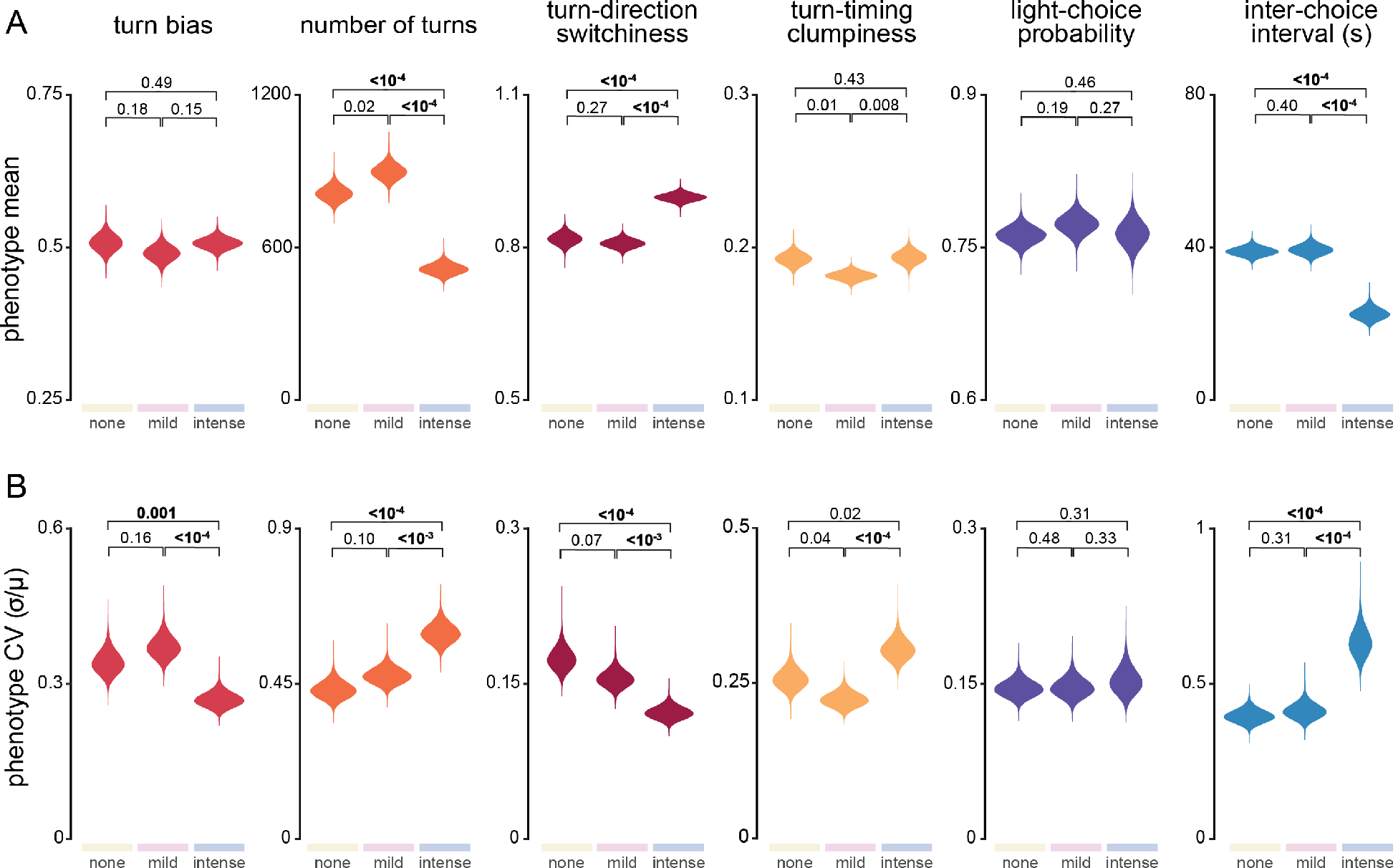
Posterior distributions of mean and intragenotypic variability (coefficient of variation; CV) for Canton-S flies under both enrichment treatments. Values shown at the top of each plot are the fraction of the posterior distribution of differences between two treatments (e.g., control and mild enrichment) that lies either below or above zero, depending on the direction of change. Given our finite posterior sampling, we cannot estimate fractions of distributions accurately below approximately 10^−4^. Bold values indicate treatments for which the 99% credible interval of the treatment effect does not include 0. Sample sizes of each experiment are provided in the Methods.

**Figure 4.**
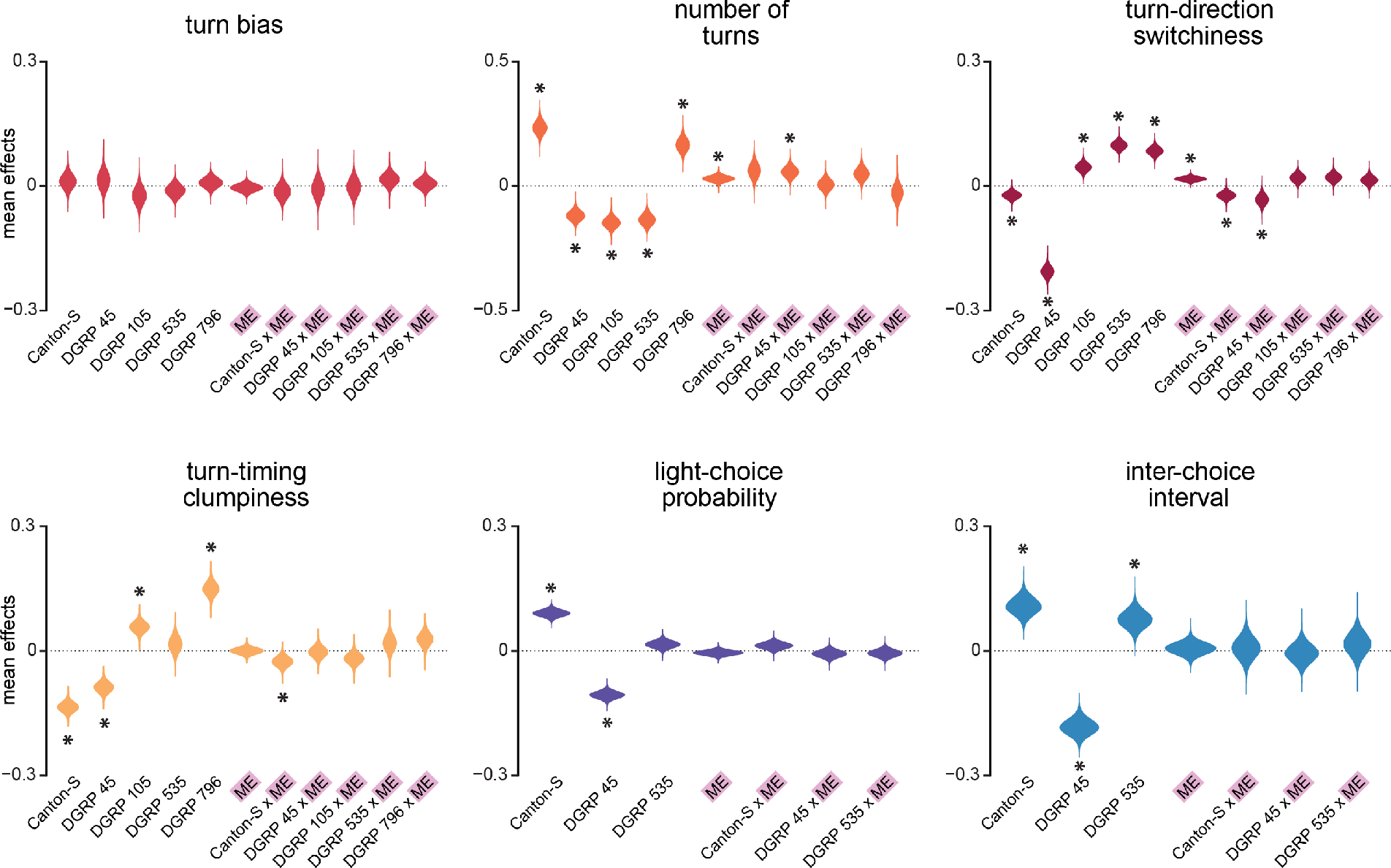
Genotype, mild enrichment, and genotype-by-mild enrichment effects on behavioral measure means. Asterisks mark those effects whose 99% credible interval does not include zero. All effects were normalized by the grand mean of all treatments and genotypes, so these values can be interpreted as effect sizes of each condition on the mean. Sample sizes of each experiment are provided in the Methods.

Mild enrichment causes small or no differences (zero effect was within the 99% credible interval) when compared to the control treatment for all the behavioral measures from both assays, with turn direction switchiness and turn timing clumpiness the most likely behavioral measures to be affected by mild enrichment. Variability in turn bias and number of turns probably increased slightly under mild enrichment, while clumpiness and switchiness probably decreased slightly. In two of these cases the direction of the effect matched the direction of the intense enrichment effect; in the other two cases, it did not. To summarize, intense enrichment had stronger effects on variability than mild enrichment, and the direction of these effects was behavior-dependent.

### The effect of mild enrichment on mean and variability is geno-type- and behavior-dependent

We were interested in testing whether the effects of enrichment we observed were dependent on genotype. We reared flies from four additional genotypes in the unenriched and mild conditions and measured their Y-maze and FlyVac behaviors. Specifically, we used four lines from the Drosophila Genetic Reference Panel (DGRP, Mackay et al., 2012). These lines (numbers 45, 105, 535 and 796) were derived from different wild-caught gravid females and then inbred for 20 generations. Thus, there is significant genetic variation between the lines, but not within them. We chose to work with these particular lines because we have previously observed that they vary in intragenotypic variability in Y-maze turn bias (Ayroles et al., 2015).

As before, we first considered effects of genotype, mild enrichment and genotype-by-mild enrichment on behavioral means (Fig. 4). Genotype had an effect (i.e., zero was not in the 99% credible interval of the posterior distribution) on all behaviors except turn bias. Mild enrichment caused a genotype-independent increase in number of turns and switchiness, but no other behaviors. There were genotype-by-mild enrichment effects on number of turns, switchiness, and clumpiness, and the directions of those effects were variable.

With respect to intragenotypic variability, we found that genotype, mild enrichment and genotype-by-mild enrichment all had effects (Fig. 5). We found that the variability of all behavioral measurements, except the number of turns, were affected by genotype. The variability of number of turns, clumpiness, and light choice increased in a genotype-independent manner under mild enrichment. Variability of switchiness and inter-choice interval were probably also increased in a genotype-independent manner by mild enrichment (a large majority of their respective posterior distributions was above zero). We observed genotype-by-mild enrichment effects for number of turns, switchiness, clumpiness, and light-choice probability. Of all the behavioral measures, switchiness showed the most variable and the strongest genotype-by-enrichment effects. To summarize, mild enrichment often increased variability in a genotype-independent fashion, but there were also frequently genotype-by-mild enrichment effects.

Interestingly, we found that the average magnitudes of mean effects were smaller than the average magnitudes of variability effects (Fig. 6A,B). For both mean and variability, genotype effects tended to be larger than the mild enrichment or genotype-by-mild enrichment effects. This pattern was especially prominent for mean effects. We also found that the sizes of the effects on behavioral means were uncorrelated with sizes of effects on variability (Fig. 6C).

**Figure 5.**
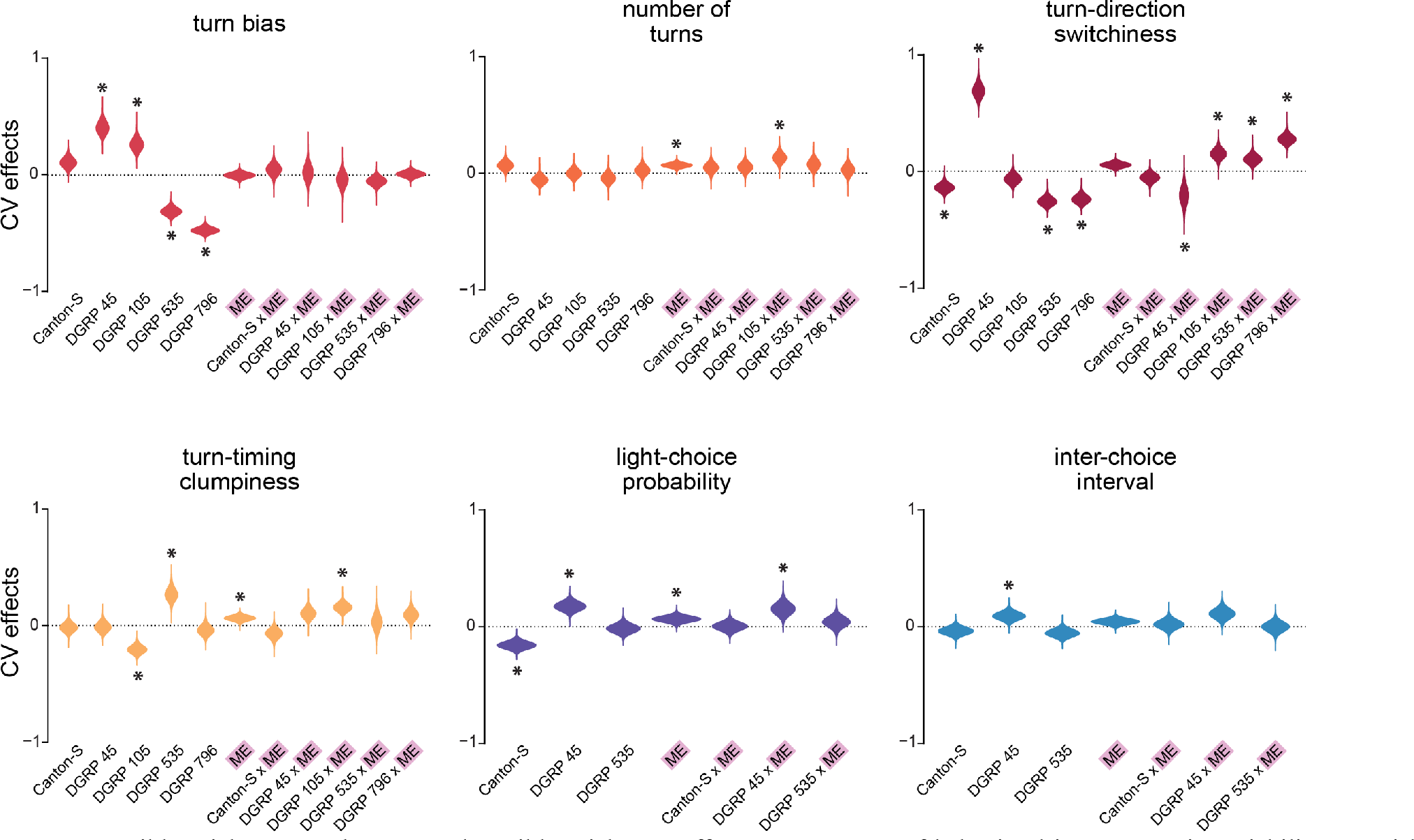
Genotype, mild enrichment, and genotype-by-mild enrichment effects on measures of behavioral intragenotypic variability. Asterisks mark those effects whose 99% credible interval does not include zero. All effects were normalized by the grand variability of all treatments and genotypes, so these values can be interpreted as effect sizes of each condition on intragenotypic variability. Sample sizes of each experiment are provided in the Methods.

## Discussion

The goal of our study was to test two opposing hypotheses about the effects of enrichment on intragenotypic behavioral variability. We hypothesized that enrichment could increase variability due to the increase in microenvironmental diversity or decrease variability due to enriched environments more closely mimicking natural conditions (resulting in more robust development of behaviors and elimination of extremes that can occur in impoverished conditions) (Körholz et al., 2018). To this end, we examined six behaviors in *Drosophila melanogaster* across several genotypes and employed two levels of enrichment. We found that for five of the six behaviors, when examined across several genotypes, mild enrichment via a fly ‘jungle gym’ likely led to an increase in intragenotypic variability, supporting the hypothesis that enrichment causes an increase in behavioral variability due to an increase in microenvironmental diversity. However, these genotype-independent effects were generally smaller than the effects of genotype or genotype-by-enrichment interactions. Therefore, the effect of enrichment on intragenotypic variability is dependent on the particular genotype and behavior being assayed. We also found that the effects of enrichment on behavioral means and variabilities were largely independent of each other, with variability effects having larger magnitudes than mean effects (Fig. 6). This finding confirms that our enrichment paradigm was able to affect both mean and variability, and that these effects are potentially independent. From our experiments, it remains uncertain which aspects of enrichment influence mean and which influence variability, or indeed, if these aspects are one and the same. Behavioral variability was also more strongly impacted by enrichment, which may underscore a biological flexibility that is not present in determining mean behavior.

With respect to both mean and variability, we found that genotype usually had a larger effect than the mild enrichment. This was especially obvious when looking at the genotype effects on behavioral means, where all behaviors except turn bias showed large genotype effects (Figs. 4,6). The lack of effect of genotype on turn bias is consistent with previous work that found no differences in the mean turn biases of 159 DGRP lines (Ayroles et al., 2015). Genotype also had strong effects on intragenotypic variability (Fig. 5), as expected (Ayroles et al., 2015; Buchanan et al., 2015; Kain et al., 2012).

We found evidence of interactions between genotype and enrichment for practically all the behavioral measures examined, though the magnitude of these interactions was behavior-dependent (Fig. 6). For example, turn bias and inter-choice interval showed very little genotype-by-enrichment effect for variability, but large effects were seen for turn switchiness. Dependence of variability on the particular parameter measured was previously noted in mouse enrichment studies (Körholz et al., 2018; Van de Weerd et al., 2002). Behavioral parameters may fall into different categories with respect to their response to enrichment. For example, switchiness is a measure of intraindividual variability (Figure 1), hinting at a link between the biological mechanism controlling variability from trial-to-trial and individual-to-individual (Stamps et al., 2013). Our results also make it clear that in assessing the effects of enrichment on a particular measure of behavior, genotype cannot be ignored. These interactions are consistent with previous findings in rats and mice (Konkle et al., 2010; Toth et al., 2011; van de Weerd et al., 1994), where the effects of enrichment differed between strains.

We also examined whether the degree of change in variability would scale with the level of enrichment presented. We raised one cohort of flies in a naturalistic setting, subject to the environmental fluctuations of the outdoors and with access to numerous organic and inorganic substrates (Fig. 2). In general, this intense enrichment had much stronger effects on our behavioral measures than the mild enrichment fly vial ‘jungle gym.’ Even though the effects of intense enrichment were more pronounced, the direction of these effects was behavior-dependent (Fig. 3). For example, we saw a decrease in intragenotypic variability for turn bias and turn switchiness under intense enrichment, but an increase in the variability for number of turns and turn clumpiness. The directions of these effects varied by behavior, even relative to the direction of the mild enrichment effect. One of our predictions was that a more naturalistic enrichment treatment could lead to a decrease in variability because of an increase in robustness, but it could also be that naturalistic enrichments cause fly populations to exhibit more natural behaviors in general, whether or not that corresponds to a decrease in behavioral variability. Future studies could address what constitutes natural fly behavior in more detail, whether by making field-deployable assays or bringing wild flies directly to lab for testing, though any comparisons with our current enrichment paradigm would need to carefully consider population genotypic variance.

**Figure 6.**
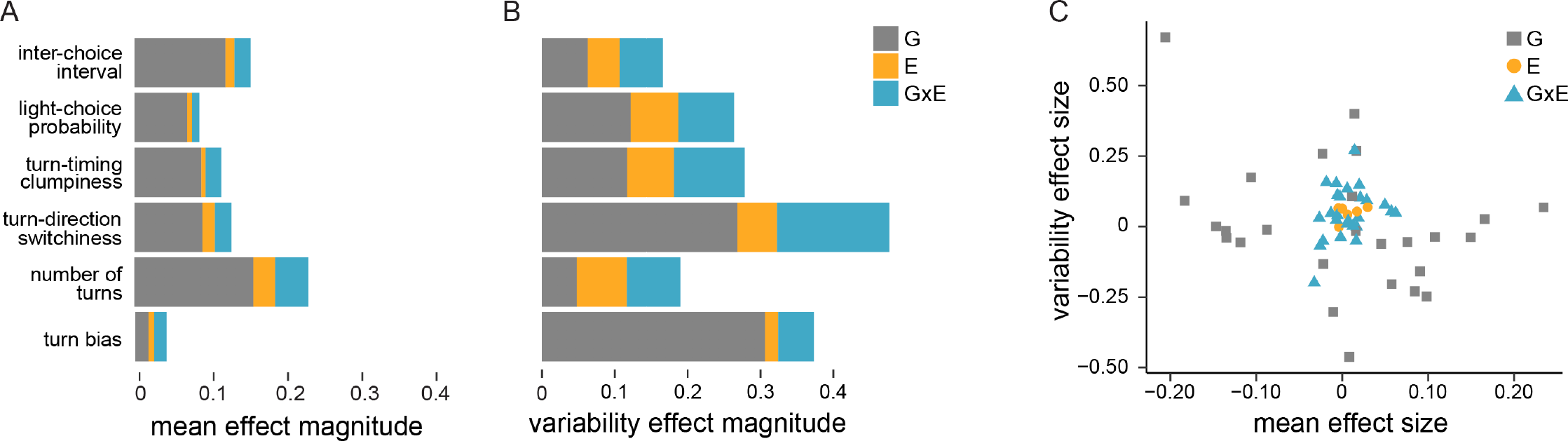
Summary of the genotype, mild enrichment, and genotype-by-mild enrichment effects on measures of behavioral mean and intragenotypic variability. A-B) Length of the bar represents the average magnitude of the effect; G is genotype effect, E is mild enrichment effect, and GxE is genotype-by-mild enrichment effect. C) Correlation of variability effect sizes and mean effect sizes across all behaviors, separated into genotype, mild enrichment, and genotype-by-mild enrichment effects. All effects were normalized by either the grand mean or variability of all treatments and genotypes, so these values can be interpreted as effect sizes of each condition.

Overall, our results support the hypothesis that enrichment increases intragenotypic variability, though this effect is dependent on the particular genotype and behavior in question. We also conclude that genotype is likely to remain the main determinant of intragenotypic variability. Our findings make it apparent that the genotype used and behavior measured will affect the inferred relationship between environmental variability and behavioral variability. Moreover, the type of enrichment (e.g., mild vs intense enrichment) can qualitatively and quantitatively alter this relationship. This, and the effects of genotype and behavioral measure, could be why effects observed in one enrichment study may not be replicated in another (Toth et al., 2011). While the multifactorial nature of enrichment provides challenges, its specific effects continue to be of great interest in behavioral research, and high throughput, data-driven approaches have the potential to illuminate the complex relationships between environmental variability and behavioral variability.

## Methods

### Data and code

All of the measures of individual behavior and all analysis scripts are publicly available at https://zenodo.org/record/2573158. They are also hosted on our lab website at http://lab.debivort.org/enrichment.

### Behavior and enrichment protocols

Fly stocks were cultured in vials on Caltech formula media at 25°C in temperature controlled incubators on a 12h-12h light-dark cycle.

For mild enrichment, in each vial of media where experimental animals were to develop, 3 female and 2 male parental flies were housed for 3-5 days. The parents were removed and the jungle gym enrichment was inserted. The enrichment object consisted of a number of craft items that were hot-glued to a wooden applicator stick that was inserted into the media. These were identically constructed for ~30 vials, with the exception of the pom-pom color, which in some vials at random was white and in the others pink. F_1_ experimental progeny developed in this enriched environment for around 10 days in incubators. Once they began eclosing, the were allowed to accumulate for 1-2 days, after which they were removed and mixed under cold anesthetization with other flies from the same genotype. They were then sorted into cohorts of 40 males and 40 females and placed in mildly enriched vials for 3-5 days. At this point, their behavior was measured in the Y-mazes for 2 hours according to the methods in Buchanan et al., 2015 (though here we loaded anesthetized experimental animals on ice to transfer them into the Y-mazes, rather than CO_2_). After the Y-maze assay, they were anesthetized on ice and returned to their mildly enriched vials for 5-10 days at which point their phototactic preferences were measured using FlyVac according to the methods in Kain et al., 2012. This mild enrichment procedure was used for flies of all genotypes.

For intense enrichment, we prepared a population cage using 1m wooden dowels to make a cubic frame, with sheer white polyester drapery material as walls. A tube of this material, normally held closed by binder clips, provided access to the inside of the cage. The items shown in Fig. 2 were introduced to the cage at the time of its construction. For the experiment, a parental Canton-S population of 200 males and 200 females was placed in the cage on September 5th 2013, and removed 9 days later. F_1_s were collected on September 28^th^, 30^th^, and October 5^th^ 2013 and assayed in the Y-mazes on those days respectively. Flies were recovered from the Y-mazes using cold anesthetization and stored in unenriched standard media tubes in groups of ~30 individuals until testing with FlyVac on October 6^th^ 2013. For all assays, males and females were tested in equal proportions.

For Y-maze enrichment experiments with Canton-S, we assayed 151 control flies, 203 mildly enriched flies, and 206 intensely enriched flies. For Canton-S FlyVac experiments, we assayed 175 control flies, 140 mildly enriched flies, and 86 intensely enriched flies.

For Y-maze enrichment experiments with the DGRP lines, we assayed:

DGRP 45: 166 control, 133 mildly enriched
DGRP 105: 130 control, 148 mildly enriched
DGRP 535: 113 control, 111 mildly enriched
DGRP 796: 132 control, 128 mildly enriched

For FlyVac enrichment experiments with DGRP lines, we assayed:

DGRP 45: 157 control, 144 mildly enriched
DGRP 535: 122 control, 140 mildly enriched

### Behavior measures and null model distributions

Behavior measures from the Y-maze assay (turn bias, number of turns, turn direction switchiness and turn timing clumpiness) were calculated from the vectors of turn directions and times that each fly produced in experiment. Behavior measures from FlyVac (light-choice probability and inter-choice interval) were calculated from the FlyVac data output file (Kain et al., 2012). These measures were computed and/or collected into a common data structure in MATLAB 2013a (The Mathworks, Inc., Natick, MA).

Null hypothesis distributions (Fig. 1) were generated in Matlab 2013a by resampling (with replacement) a million values for each distribution as follows: 1) For turn bias and light-choice probability, a) all observed choice values (i.e. left vs. right and light vs. dark) were pooled across individuals, b) an individual was chosen at random from all tested, and a vector of length equal to the number of behavioral choices performed by that individual during the experiment was populated randomly with values from the pool, and c) the turn bias or light-choice probability for that vector was recorded. 2) For number of turns, a) the observed inter-turn intervals (ITIs) were pooled across individuals, b) ITIs were chosen randomly one at a time until their cumulative sum exceeded 7200000ms, the length of an experiment, and c) the number of turns in this sequence was recorded. The moderate discrepancy in mean between the null hypothesis distribution and experimental distribution in this analysis arises from the disproportionate number of short intervals contributed to the total pool by more active animals. 3) For switchiness, which arises from slight dependence between consecutive turns in the LR turn sequence (Ayroles et al., 2015), a) we implemented a Markov chain in which the L-L (= R-R) transition probabilities yielded LR sequences with mutual information between successive turns equal to the observed mutual information (0.018 bits, P(L-L) = 0.592). b) An individual was chosen at random from all tested, and a choice sequence of length equal to the number of choices performed by that individual during the experiment was generated using the Markov chain, and c) the switchiness of this sequence (= (#LR+#RL)/(2*turn bias*(1-turn bias)*num turns) was recorded. 4) For clumpiness, a) the observed ITIs were pooled across individuals, b) an individual was chosen at random from all tested, and a vector of length equal to the number of choices performed by that individual during the experiment was populated randomly with values from the ITI pool, and c) clumpiness (= MAD(ITIs)/(sum(ITIs)/num turns), where MAD is the median absolute deviation from the median) was recorded. Thus, this approach reflects sampling variation in number of turns and mean ITI as well as clumpiness. 5) For inter-choice interval (ICI), a) the observed ICIs were pooled across individuals, b) an individual was chosen at random from all tested, and a vector of length equal to the number of choices performed by that individual during the experiment was populated randomly with values from the ICI pool, c) the mean ICI across this vector was recorded.

### Bayesian inference of mean and variance effects

To get the estimates of the posterior distributions of behavioral mean and variance, and the effects on the observed distributions of enrichment treatment, genotype, and their interactions, we employed linear and generalized linear models in R’s Stan interface v.2.18.2 (The Stan Development Team). The Stan platform allows the user to specify desired models and performs full Bayesian inference using Hamiltonian Monte Carlo with the No U-Turn sampler (Carpenter et al., 2017). To get the posterior distributions of the mean and variance for turn bias, light-choice probability, switchiness, and clumpiness under different enrichment-genotype conditions, we specified the following model:

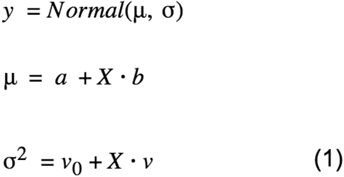

where *y* is a vector of the behavioral outcomes of individuals that comes from a normal distribution with parameters (mean) and (standard deviation). *μ* and *σ*^2^ are specified via linear models, where *a* and *v*_0_ are intercepts, *X* is a logical predictor matrix specifying the genotype and/or enrichment treatment for each individual, and *b* and *v* are vectors of coefficients of the linear model.

Since the distribution of the number of turns was right-skewed, bounded to real positive integers, and over-dispersed compared to a Poisson distribution, we chose to model this measure with a negative binomial as follows:

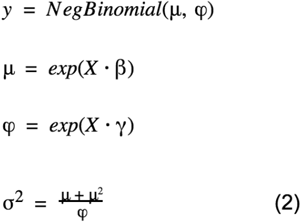

Here, *y* is the vector of number of turns of individuals modeled by a negative binomial distribution with parameters *μ* (mean) and *σ* (dispersion). Both parameters are related to the coefficients of a generalized linear model (*β* and *γ*) via a log-link function. *X* is the experimental design matrix, as above. *σ*^2^ is the variance calculated from the mean and dispersion parameters.

To model inter-choice intervals, we chose a gamma distribution since the data is right-skewed and positive continuous:

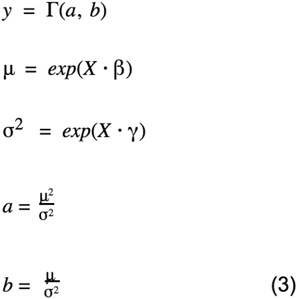

Where *a* and *b* are the shape and rate parameters of the gamma distribution, respectively, and the rest of the parameters are as above.

For the estimation of all posterior distributions, we set our priors on the coefficients to broad Cauchy distributions centered at zero to allow them to be weakly informative (Gelman et al., 2008). Our qualitative findings were robust to the choice of prior, as uninformative uniform told the same story (Fig. S3). To sample posteriors, we used four chains and 50,000 - 100,000 iterations per chain, with the target average proposal acceptance probability of 0.8-0.9 and maximum tree depth of 10-15, to generate a posterior distribution of 100,000 - 200,000 samples (50% of chain iterations were used for tuning the Hamiltonian Monte Carlo sampler parameters and were discarded as the burn-in period). The ratio of chain effective sample size to sample size was in the range of 0.7 - 1.2, indicating that posterior estimate error due to autocorrelation was minimal. To get posterior distributions for the coefficients of variation, we took the square root of the variance and divided it by the mean at each step in the chain. To check our model fits, we carried out graphical posterior predictive checks (Fig. S4), and found that our models fit the data well.

We adapted the methodology used by Kruschke’s BEST method (Kruschke, 2013) to our own posterior distributions in order to determine which effects were inferred to differ from zero. To estimate the posterior distribution of a treatment effect, we subtracted the parameter values of one treatment condition from the control (or the other treatment condition) at each step in the chain and took the distribution of that difference. We calculated the 99% highest density interval (the credible interval) of the posterior of treatment effects to evaluate whether the treatments had an effect - if the 99% highest density interval excluded 0, we inferred an effect between the treatments. Since this approach is subject to multiple comparisons concerns, we chose the 99% credible interval (rather than e.g., 95%) as more stringent indicator of effects. While we think that the 99% credible interval is a useful guide to pulling out the strongest effects we observe, we believe that the strength of Bayesian inference lies in being able to examine the posterior distributions as a whole and observing their trends (rather than applying a threshold to identify effects).

To determine the contribution of genotype, mild enrichment, and genotype-by-mild enrichment effects to the variability (coefficient of variance) and mean of each behavior, we used the following formulas:

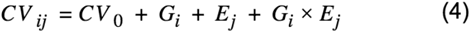

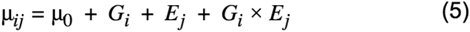

*CV*_*ij*_ and *μ*_*ij*_ are the variability and mean, respectively, of genotype *i* in treatment group *j*. *CV*_0_ and *μ*_0_ are the grand variability and mean, averaged over all genotypes and treatments. *G*_*i*_ is the deviation of the variability or mean of genotype *i* from the grand parameter in question, calculated over all treatment groups. *E*_*j*_ is the deviation of treatment group *j* from the grand parameter, calculated over all genotypes. The treatment groups in this experiment were mild enrichment or control vials. *G*_*i*_ × *E*_*j*_ is the specific deviation of genotype *i* in treatment group *j* after accounting for the main effects of genotype *i* and treatment group *j*. All deviations were standardized by dividing them by the grand parameter value in order to interpret them as effect sizes.

## Acknowledgements

We are grateful to Kyle Honegger for discussions of Bayesian inference and Nico O. Wagner for feedback on the manuscript. We also thank Carolyn Elya for moral support when the sampler misbehaved. SH was supported by the Harvard PRISE program. JA was supported by the National Science Foundation PoLS-HF-SRN grant (NSF-1806818). BdB was supported by a Sloan Research Fellowship, a Klingenstein-Simons Fellowship Award, a Smith Family Odyssey Award, and the National Science Foundation under grant no. IOS-1557913.

## Conflict of Interest

The authors declare no competing interests.

## Supplementary figures

**Figure S1.**
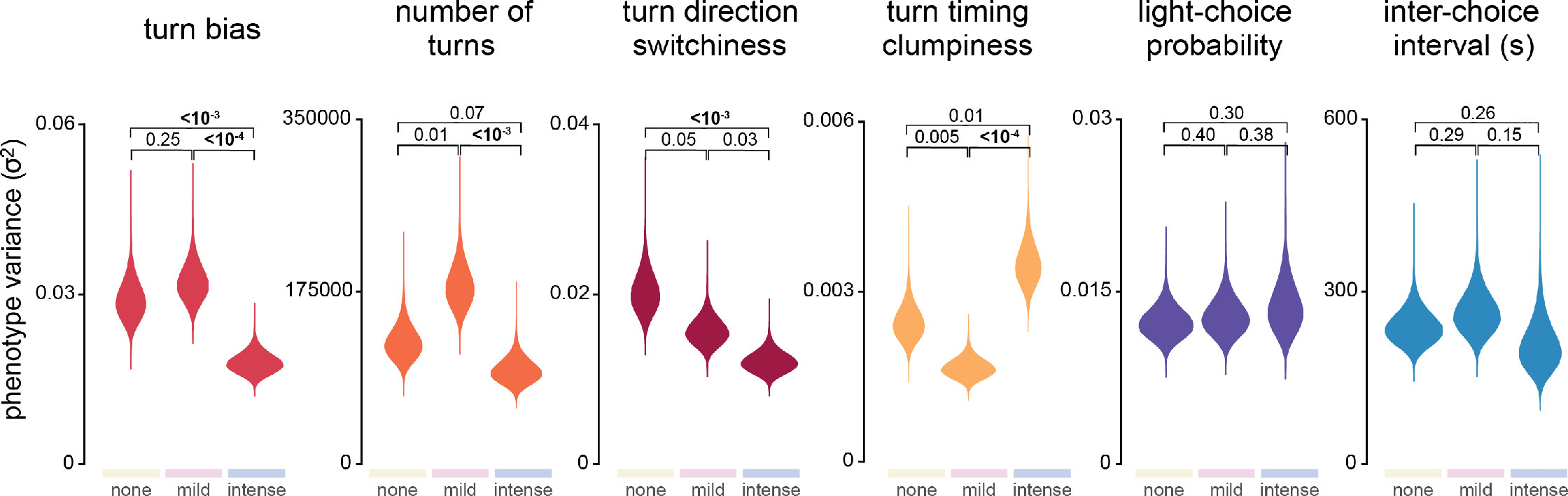
Posterior distributions of behavioral measure variance for Canton-S flies under two enrichment treatments. Numbers at the tops of the panels are the fraction of the posterior distribution of differences between two treatments (e.g., control and mild enrichment) that lies either below or above zero, depending on the direction of change. Given our finite posterior sampling, we cannot accurately estimate fractions of distributions below approximately 10^−4^. Bold values indicate treatments for which the 99% credible interval of the treatment effect does not include 0. See Methods. Sample sizes of each experiment are provided in the Methods.

**Figure S2.**
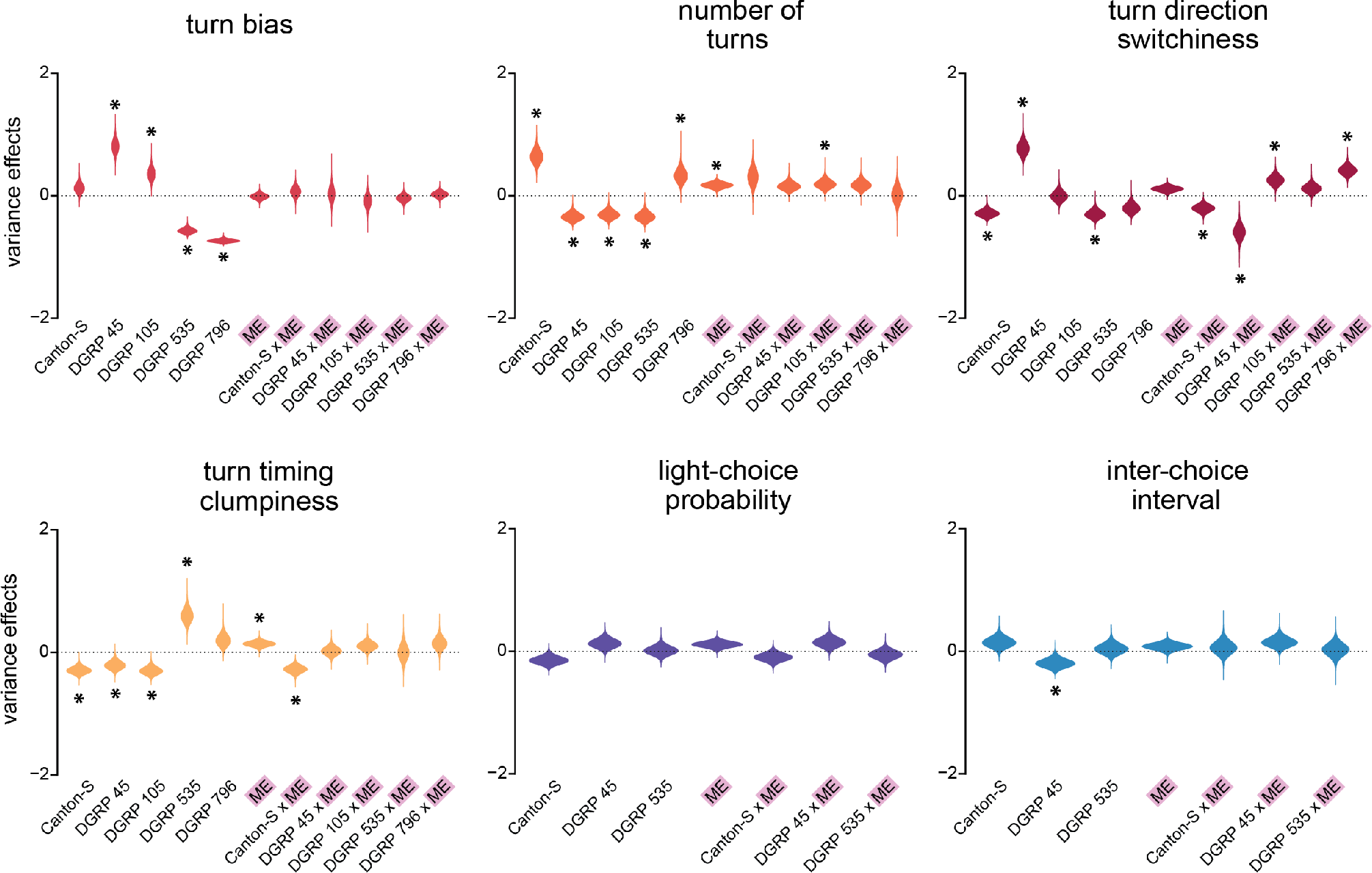
Genotype, mild enrichment, and genotype-by-mild enrichment effects on behavioral metric variance. Asterisks mark those effects whose 99% credible interval does not include zero. All effects were normalized by the grand variance of all treatments and genotypes. Sample sizes of each experiment are provided in the Methods.

**Figure S3.**
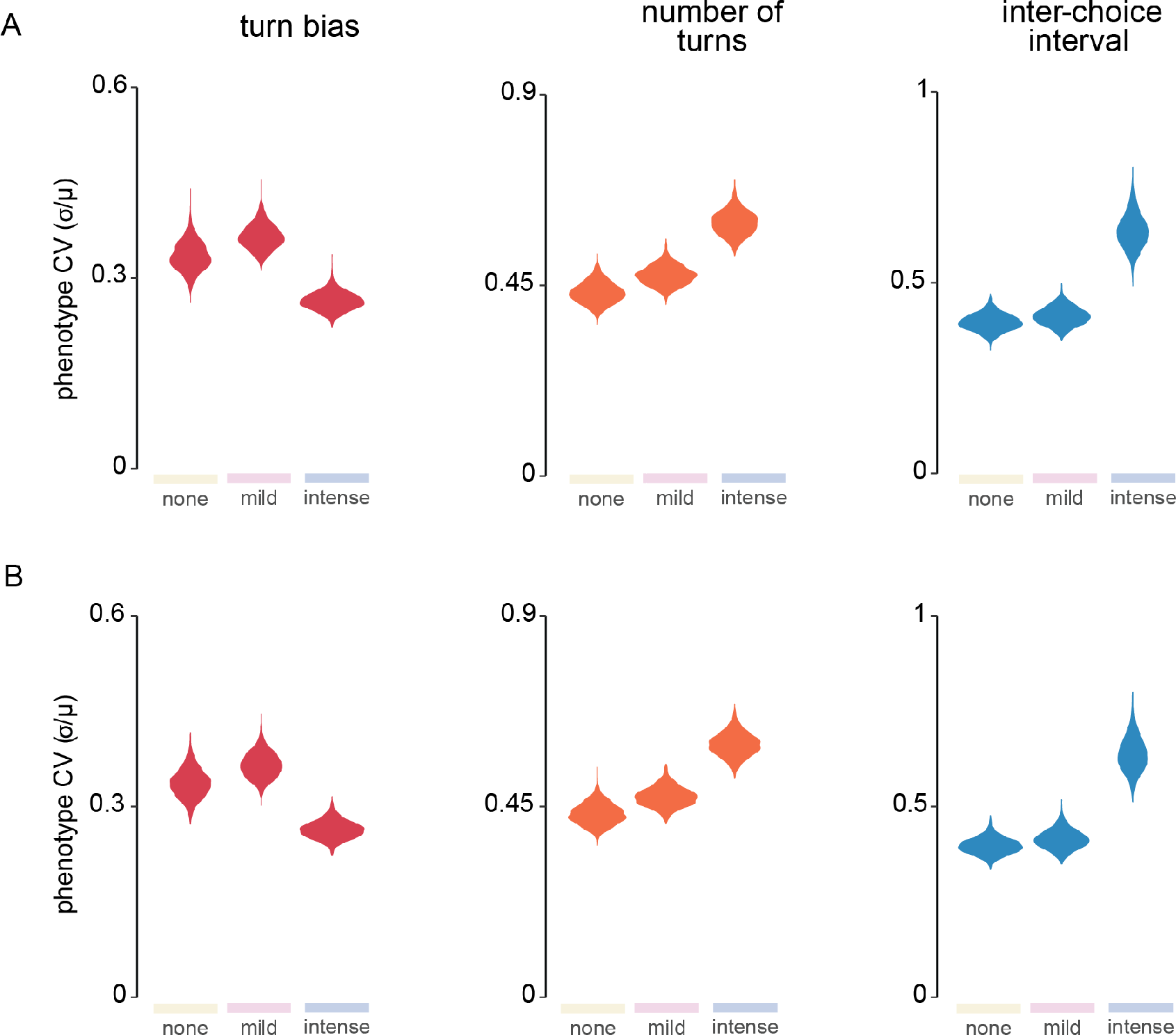
Weakly informative cauchy and uniform priors result in similar posterior distributions. A) Posterior distributions generated under a Cauchy prior. B) Posterior distributions generated under a uniform prior. Thus, our choice of a weakly informative prior, rather than non-informative prior, is not driving our conclusions about the treatment effects. Turn bias, number of turns, and inter-choice interval were chosen to represent the three types of models used: normal, negative binomial, and gamma, respectively. Sample sizes of each experiment are provided in the Methods.

**Figure S4.**
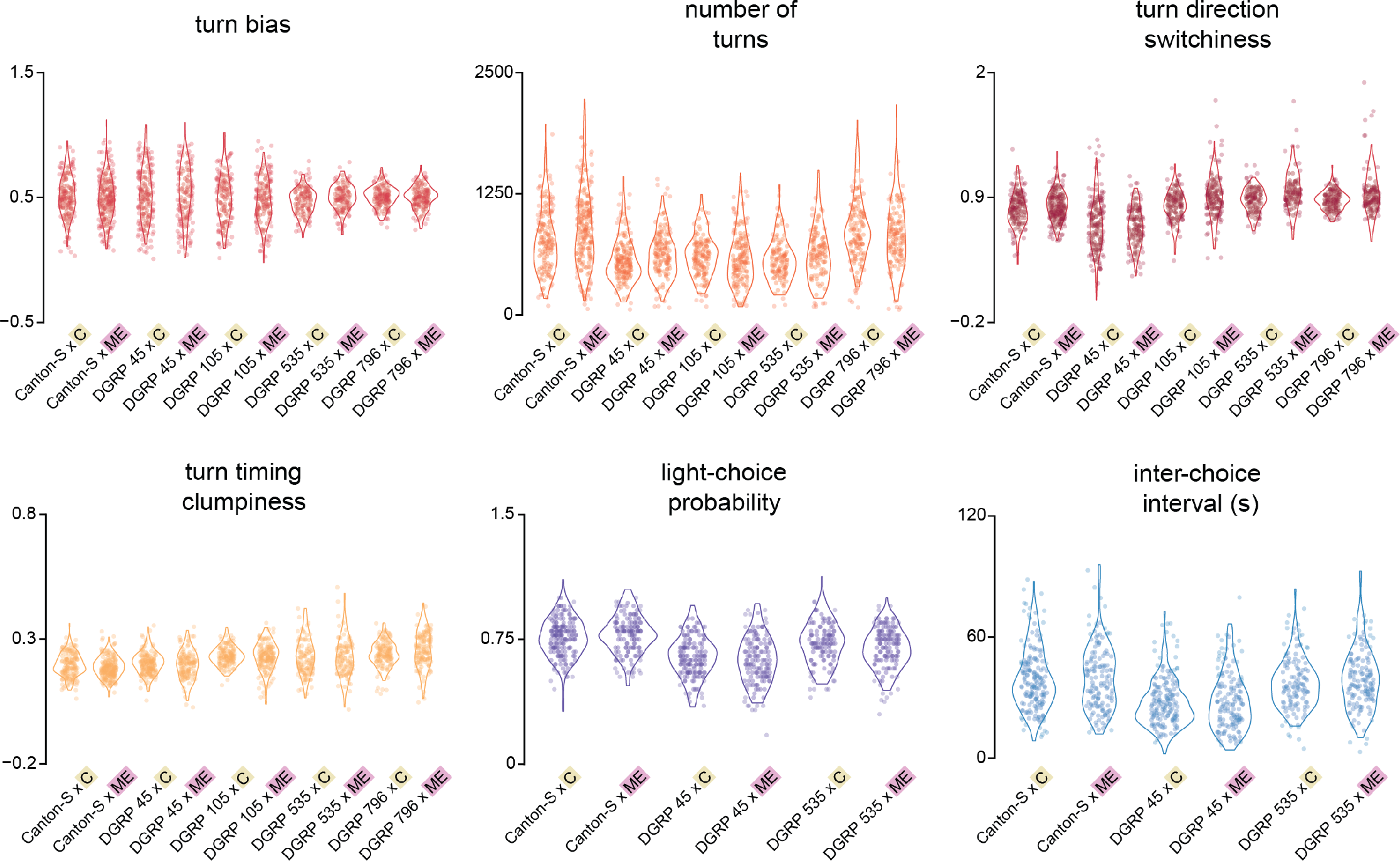
Concordance between values sampled from our models of treatment effect parameters and the experimental data (posterior predictive checks). Violin plots are kernel density estimates from a sample of values drawn from distributions whose parameters are drawn from our effect posteriors. Points are the original data. The general agreement between the modeled and empirical distributions indicates that we have appropriately modeled the effects and that our choice of priors was not wholly inappropriate. Sample sizes of each experiment are provided in the Methods.

